# How Much Information is Provided by Human Epigenomic Data? An Evolutionary View

**DOI:** 10.1101/317719

**Authors:** Brad Gulko, Adam Siepel

## Abstract

Here, we ask the question, “How much information do available epigenomic data sets provide about human genomic function, individually or in combination?” We consider nine epigenomic and annotation features across 115 cell types and measure genomic function by using signatures of natural selection as a proxy. We measure information as the reduction in entropy under a probabilistic evolutionary model that describes genetic variation across ∼50 diverse humans and several nonhuman primates. We find that several genomic features yield more information in combination than they do individually, with DNase-seq displaying particularly strong synergy. Most of the entropy in human genetic variation, by far, reflects mutation and neutral drift; the genome-wide reduction in entropy due to selection is equivalent to only a small fraction of the storage requirements of a single human genome. Based on this framework, we produce cell-type-specific maps of the probability that a mutation at each nucleotide will have fitness consequences (*FitCons* scores). These scores are predictive of known functional elements and disease-associated variants, they reveal relationships among cell types, and they suggest that ∼8% of nucleotide sites are constrained by natural selection.

## INTRODUCTION

Recent technological advances have enabled the generation of massive quantities of genomic data describing natural genetic variation as well as diverse epigenomic features such as chromatin accessibility, histone modifications, transcription factor binding, DNA methylation, and RNA expression^1-5^. However, the capability to gain insight into key cellular functions from this noisy, high-dimensional data has considerably lagged behind the capacity for data generation. Indeed, while the available data allows the vast majority of the human^1^ and mouse^2^ genomes to be associated with some type of “biochemical function”, often in a cell-type-specific fashion, it is unclear—and highly controversial^6-9^—to what degree this biochemical function reflects critical roles in cellular processes that have bearing on evolutionary fitness, as opposed to representing, say, noisy or incidental chromatin accessibility, protein/DNA binding, or transcription. This uncertainty about the true biological significance of many high-throughput epigenomic measurements is a critical barrier not only for interpretation of the available data, but also for prospective decisions about how much new data to collect, of what type, and in what combinations.

Numerous computational methods have recently been developed to address the problem of extracting biological meaning from large, heterogeneous collections of high-throughput genomic data by a variety of different strategies. These include methods that cluster genomic sites based on epigenomic patterns^10-12^, machine-learning predictors of pathogenic variants^13,14^ or molecular phenotypes^15-17^, and methods that combine epigenomic data with patterns of polymorphism or cross-species divergence to identify regions under evolutionary constraint^18-20^. Our contribution to this literature has been to develop a probabilistic framework, called *INSIGHT* ^21,22^, for measuring the bulk influence of natural selection from patterns of polymorphism and divergence at collections of target sites, and methods that combine this framework with epigenomic data to estimate the probability that a mutation at any position in the genome will have fitness consequences^23,24^. Our predictors of fitness consequences, called *FitCons* and *LINSIGHT*, are competitive with the best available methods for predicting known regulatory elements and disease-associated variants, but, importantly, they also provide insight into how much of the genome is influenced by natural selection and the distributions of fitness effects at selected sites. Nevertheless, neither our methods, nor other available methods, directly address questions about the global information associated with epigenomic data, such as, “How much did data set *X* reveal about genomic function?” or “How much new information will experiment *Y* provide?”

Here, we adapt our evolution-based *INSIGHT* framework to directly address the question of how much information about genomic function is provided by general epigenomic “features,” including both genome annotations and high-throughput epigenomic data sets. The premise of our approach is that signatures of natural selection in DNA sequences can serve as a proxy for genomic function by reflecting fitness constraints imposed by cellular functions. We develop a novel information-theoretic framework for simultaneously clustering genomic sites by combinations of epigenomic features and evaluating the strength of natural selection on these sites. In addition to allowing us to evaluate relative amounts of global information provided by these epigenomic features, individually and in combination, this approach produces a collection of 115 cell-type-specific genome-wide maps of probabilities that mutations at individual nucleotides have fitness consequences (*FitCons* maps), which we demonstrate are richly informative. Together, our analyses not only provide a guide for data interpretation and experimental design, but they also shed light on the fundamental manner in which biological information is stored in the genome.

## RESULTS

### *FitCons2* finds a clustering of genomic sites that maximizes information about natural selection

The main idea behind the *FitCons2* algorithm is to recursively partition sets of genomic sites into two subsets, according to their associated epigenomic and annotation features (Figure 1). For example, at a particular step the algorithm might subdivide a given set of genomic sites, itself defined by a previous partitioning, into sites showing high and low transcriptional activity, based on counts of aligned RNA-seq reads, or those showing high and low chromatin accessibility, based on DNase-seq data. At each step in the algorithm, all candidate partitions of all currently active sets are considered, based on a collection of pre-discretized data types (Figure 1a&b). The algorithm selects the decision rule for partitioning that most improves the goodness of fit of the *INSIGHT* evolutionary model to the set of genomic sites under consideration when the model is fitted separately to the two proposed subsets rather than once to the entire set (see Methods). The procedure terminates when no partition improves the fit of the model by more than a predefined threshold (Figure 1c). In this way, the recursive algorithm produces a *K*-leaf binary decision tree that applies to each genomic site, causing the site to be assigned one of *K* labels based on its combination of local features (Figure 1d). When applied to all sites, the algorithm defines *K* clusters of genomic sites that reflect the natural correlation structure of both the functional genomic and the population genomic data. Importantly, the identified clusters tend to be homogeneous but distinct from one other in terms of their influence from natural selection on the relatively recent time scales measured by *INSIGHT*, based on both human polymorphism and divergence with nonhuman primates. The overall influence of natural selection on each cluster is summarized by the associated estimate of the *INSIGHT* parameter *ρ,* which can be interpreted as the probability that a point mutation will have fitness consequences. As in the original *FitCons* algorithm, these *ρ* estimates are mapped back to the corresponding genomic sites and treated as nucleotide-specific fitness-consequence (FitCons) scores (Figure 1e).

**Figure 1.**
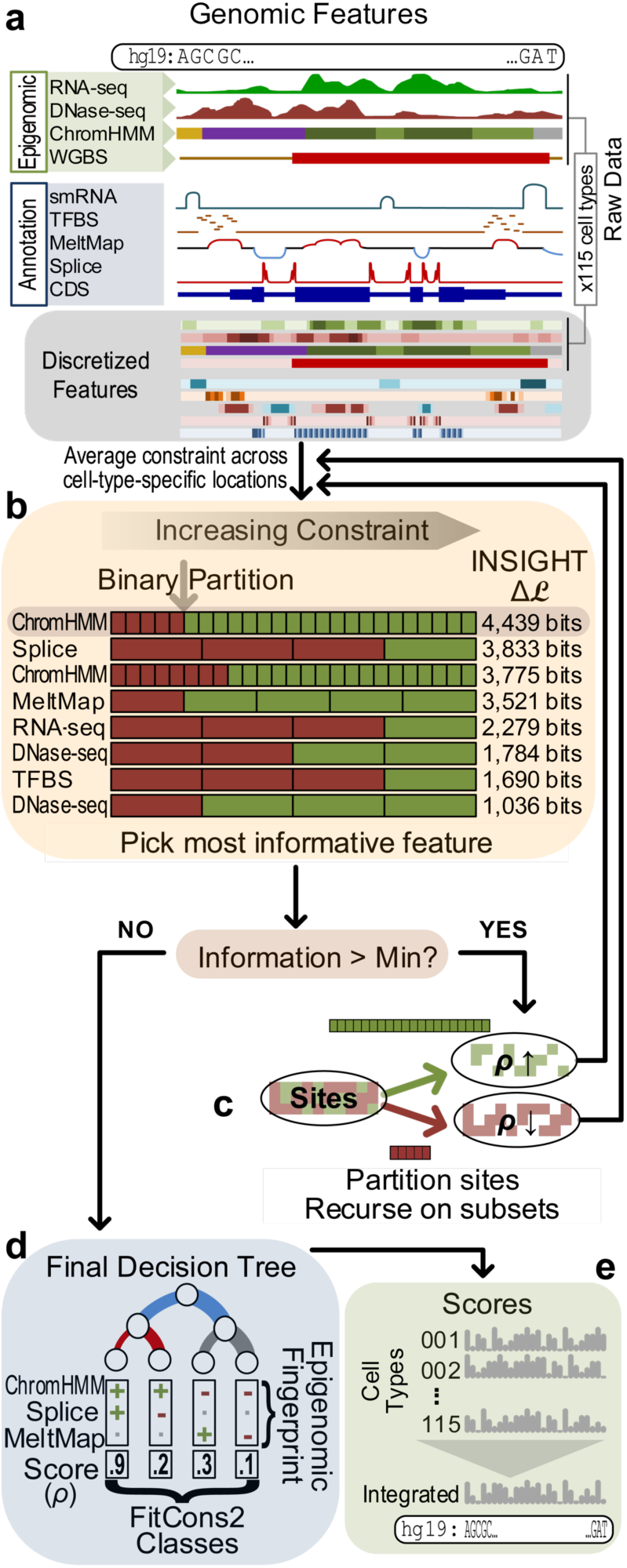
Conceptual diagram of *FitCons2* algorithm. (a) The algorithm starts with raw data describing genomic features along the human genome sequence (assembly hg19). Two types of features are considered: four Epigenomic features, represented separately for each of 115 cell types, and five Annotations, shared across cell types (see Table 1). In pre-processing, the raw data sets are discretized into between 2 and 25 feature classes, which are ordered by their estimated *ρ* values (Methods and Table 1). (b) The algorithm builds a decision tree by recursively finding binary partitions of “active sets” of genomic positions (corresponding to leaves in the growing tree). At each step, it considers all possible binary partitions of each active set. Each binary partition is defined by applying a threshold to an ordered, discretized feature (gray arrow). The algorithm selects the active set (leaf) and binary partition that are maximally informative about selection. Information is measured by the increase in log likelihood (Δ*ℒ*) obtained by fitting the *INSIGHT* model separately to each subset of genomic sites rather than once to the entire active set (Methods). During this procedure, the algorithm averages over cell-type-specific locations for the Epigenomic features. (c) The recursive process is repeated until the improvement in information fails to exceed a minimal threshold. (d) The end result is a *K*-leaf decision tree such that each internal node represents a binary decision rule and each leaf corresponds to a combination of decision rules that can be applied to each nucleotide site in the genome. Each of these *K* combinations of decision rules induces a cluster of genomic sites that share a particular epigenomic “fingerprint”. Each of these *K* clusters is also associated with an estimate of *ρ* from *INSIGHT* (its *FitCons2* score). (e) These estimates of *ρ* can be mapped back to the genome sequence separately for each cell type. An “integrated” score that summarizes all cell types is also computed (Methods).

When evaluating candidate decision rules, the *FitCons2* algorithm measures the goodness of fit of the *INSIGHT* model in terms of its log likelihood. The negated log likelihood, however, can be viewed as an estimate of the *entropy* of the probability distribution induced by the model, which in this case can in turn be viewed as a measure of the genetic entropy in a human population, generated and maintained since our divergence from our non-human primate ancestors (see Discussion and Methods). Therefore, the increase in log likelihood associated with a decision rule in the *FitCons2* algorithm can be interpreted as a reduction in entropy, or equivalently, as a measure of the *information gain* associated with the corresponding bi-partition of genomic sites. This quantity can further be interpreted as an indirect measure of information about genomic function in humans, under the assumption that the tendency for natural selection to constrain genetic variation is a reasonable proxy for the functional importance of DNA sequences. Thus, the recursive bi-partitioning algorithm finds a clustering of genomic sites that maximizes information about natural selection in humans, and approximately maximizes information about genomic function. Moreover, as explored below, the algorithm naturally provides a quantitative measure of how much information is contributed by individual or combined decision rules or feature types.

### An analysis of Epigenomic Roadmap data yields an informative decision tree and fitness-consequence maps for 115 cell types

We applied *FitCons2* to functional genomic data for human cells from the Roadmap Epigenomic Project^5^, together with population genomic data previously compiled for *INSIGHT* ^22-24^, consisting of polymorphism data from 54 unrelated humans and phylogenetic divergence data from alignments of the chimpanzee, orangutan and rhesus macaque genomes to the human reference genome. (Larger data sets of human polymorphism data are now available but have negligible impact in this setting; see Methods.) To summarize the Roadmap data and associated genomic annotations, we made use of nine feature types spanning a broad range of biological processes, levels of genomic resolution, and degrees of cell-type specificity, including RNA-seq, DNase-seq, small RNAs (smRNA), chromatin states (ChromHMM), annotated coding sequences (CDS) and splice sites (Splice), transcription factor binding sites (TFBS), and predicted DNA melting temperatures (MeltMap) (see Table 1). The cell-type-specific “Epigenomic” features were collected separately for each of the 115 karyotype-normal cell types represented in the Roadmap Epigenomic Project.

**Table 1:** Summary of Epigenomic and Annotation Features Used by *FitCons2*

| Name | Description | Type<sup>1</sup> | Levels<sup>2</sup> | Source |
| --- | --- | --- | --- | --- |
| CDS | Coding sequences | Annotation | 5: Start, Codon pos. 1,2,3, Non | GENCODE<sup>46</sup> |
| Splice | Splice sites | Annotation | 4: Core, Prox, Dist, Non | GENCODE<sup>46</sup> |
| MeltMap<sup>3</sup> | Predicted DNA melting temperature | Annotation | 5: VHi, Hi, Med, Lo, VLo | ref. <sup>49</sup> |
| TFBS | Transcription factor binding sites | Annotation<sup>4</sup> | 4: Hi, Med, Lo, None<sup>5</sup> | Ensembl<sup>50</sup> & ref. 22 |
| smRNA | Small RNAs | Annotation<sup>4</sup> | 4: Hi, Med, Lo, None | ENCODE & Human Epigenome Atlas<sup>6</sup>. |
| RNA-seq | Transcription | Epigenomic | 4: Hi, Med, Lo, None | Roadmap |
| DNase-seq | Chromatin accessibility | Epigenomic | 4: Hi, Med, Lo, None | Roadmap |
| ChromHMM<sup>7</sup> | Chromatin modifications | Epigenomic | 25: (see ref. 5) | Roadmap |
| WGBS | DNA methylation | Epigenomic | 2: Hypo, non-Hypo | Roadmap |

<sup>1</sup> Annotations are shared across all cell types, whereas Epigenomic data sets are specific to each cell type (115 instances of each).
<sup>2</sup> Number of discrete levels followed by labels for levels. Features that had no natural ordering (CDS, Splice, MeltMap, TFBS, ChromHMM) were ordered by estimates of $\rho$ (see Methods).
<sup>3</sup> Predicted DNA melting temperature (MeltMap) is highly correlated with G+C content on a global level but carries additional local information. Predictions depend on the DNA sequence only and are therefore considered “Annotations”.
<sup>4</sup> Owing to sparse data, the TFBS and smRNA features were based on data pooled across cell types and therefore were treated as Annotations rather than as cell-type-specific Epigenomic data (see Methods)
<sup>5</sup> Grouped by information content of motif position (see Methods)
<sup>6</sup> Based on ENCODE cell types CD20 and HUVEC, and Human Epigenome Atlas V9 samples BGM (Brain Germinal Matrix), UCSF4 embryonic stem cells, and PFK (Penis Foreskin Keratinocyte).
<sup>7</sup> Based on the 25-state version of the Roadmap ChromHMM model, which makes use of 11 histone marks and DNase-seq data (imputed where necessary).

The recursive *FitCons2* algorithm was applied to these data as described above, except that a single decision tree was estimated by averaging across all cell types when evaluating candidate decision rules (see Methods). The algorithm identified 61 classes, each defined by a distinct combination of epigenetic features and selective pressure (Figure 2 and Supplementary Tables 1 & 2). Importantly, while each of these 61 classes is associated with a single estimate of *ρ* (representing its *FitCons2* score), each class corresponds to a different set of genomic sites in each cell type, owing to differences in the cell-type-specific features. Therefore, when these class-specific estimates of *ρ* are mapped to the genome, 115 cell-type-specific *FitCons2* maps are obtained. (These maps are available as genome-browser tracks at http://compgen.cshl.edu/fitCons2/.)

**Figure 2.**
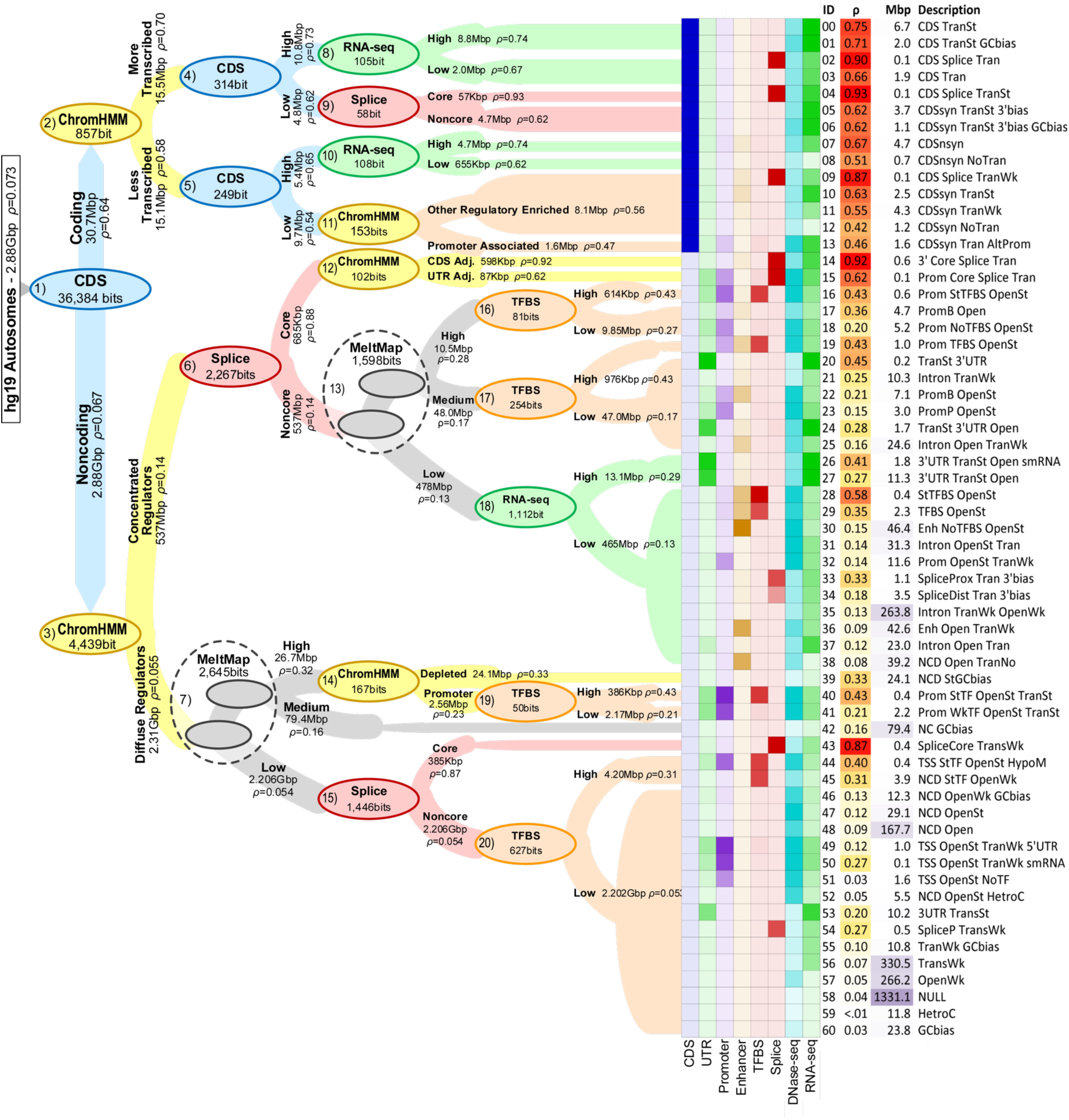
Decision tree and clusters for the human genome. The decision tree obtained by applying *FitCons2* to the human genome sequence (hg19 assembly; autosomes only). Nodes (ovals) represent decision rules (bi-partitions) and are labeled with the feature on which each rule is based as well as the associated increase in information (in bits). Nodes are colored by feature type. Edges extending from parent nodes to children are labeled with descriptions of partitions, their sizes in millions of basepairs (Mbp), and corresponding estimates of *ρ*. Edge widths are proportional to log(size). Dashed circles indicate successive binary partitions based on MeltMap that effectively create three-way splits. For simplicity, only the first 4–5 levels of the tree are shown in detail. The 61 leaves of the tree (at right) are labeled by unique identifiers, estimates of *ρ*, sizes in Mbp, and brief descriptions of the associated clusters (see Supplementary Tables 1 & 2 for additional details). Heatmap to left of cluster IDs displays relative enrichments for several annotations (Coding Sequences [CDS], Untranslated Regions [UTRs], Promoters, Enhancers, annotated Transcription Factor Binding Sites [TFBS], Splice sites, DNase-seq, and RNA-seq data).

The decision tree estimated by *FitCons2* (Figure 2) is richly descriptive about the distribution of evolutionarily relevant information across the human genome. Not surprisingly, the first partition the algorithm selects (node #1 in Figure 2) is between about 31 Mbp of annotated protein-coding sequences (CDS; *ρ* = 0.641) and the remaining noncoding sites (*ρ* = 0.067) in the genome. The partitions at the second level are based on ChromHMM states. In noncoding regions, the second split (node #3) is between a diverse collection of 20 chromatin states that are associated with regulatory and transcriptional activity (*ρ* = 0.14) and the remaining five states associated with repression, quiescence, or heterochromatin (*ρ* = 0.055). In coding regions, the second split (node #2) is between chromatin states that are associated with active transcription (*ρ* = 0.70), and ones that are not (*ρ* = 0.58).

Beginning at the third level in the tree, the partitions become highly dependent on previous splits. For example, in noncoding regions labeled with regulation-associated chromatin states, which tend to fall near exons, the next split (node #6) distinguishes a small set of nucleotides (685 kbp) that are associated with splicing and display exceptionally strong constraint (*ρ* = 0.88) from the remaining nucleotides (*ρ* = 0.14). Subsequent splits in this subtree make use of MeltMap (node #13), RNA-seq (node #18), chromatin states that identify CDS-and UTR-adjacent sites (node #12), and annotated TFBSs (nodes #16 & #17). Some similar patterns are observed outside concentrated regulators in noncoding regions, but here MeltMap is used earlier (node #7; in part as a guide to promoters and UTRs), splice sites show up later (node #15), chromatin states are used to identify promoters (node #14), and RNA-seq does not appear, presumably because these regions tend to be farther from exon boundaries. Interestingly, TFBSs are particularly informative in combination with promoter-associated chromatin states (nodes #14 & #19), which signal cell-type-specific activity. In coding regions, the third level in the tree distinguishes “high information” positions (such as start codons and 1st and/or 2nd codon positions) from “low information” positions (nodes #4 & #5). The subsequent splits in this subtree make use of information such as RNA-seq and overlap with splice sites, all features that are naturally more informative inside coding regions than outside. Notably, exons with high RNA-seq signal are found to be substantially more constrained than those with lower RNA-seq signal (nodes #8 & #10), consistent with previous findings^25^. Altogether, *FitCons2* identifies a diverse collection of clusters in the genome, ranging in size from very small (∼60 kbp) to very large (the “NULL” class [58] accounts for over 1 Gbp), and with *ρ* values from <1% to 93%. The information-based algorithm naturally finds tradeoff between large clusters with weak signals of constraint and small clusters with strong signals of constraint, both of which can produce large aggregate reductions in entropy, and it also discovers highly informative combinations of features.

### A few genomic features contribute most of the information about selection

The reduction in entropy across the entire decision tree, measured at 58,759 bits, can be interpreted as the total information about selection (and, indirectly, about genomic function) provided by all of the available functional genomic data and annotations. Moreover, feature-specific contributions to this total can be obtained by summing over all nodes (decision rules) that make use of each feature. These estimates (Figure 3a, orange bars) suggest that 62.9% of all of the available information is attributable to CDS annotations (as reported above), followed by 11.0% from ChromHMM, 8.4% from MeltMap, 7.2% from Splice, 4.8% from RNA-seq, 2.9% from TFBSs, and < 2% from each of DNase-seq, WGBS, and smRNA.

**Figure 3.**
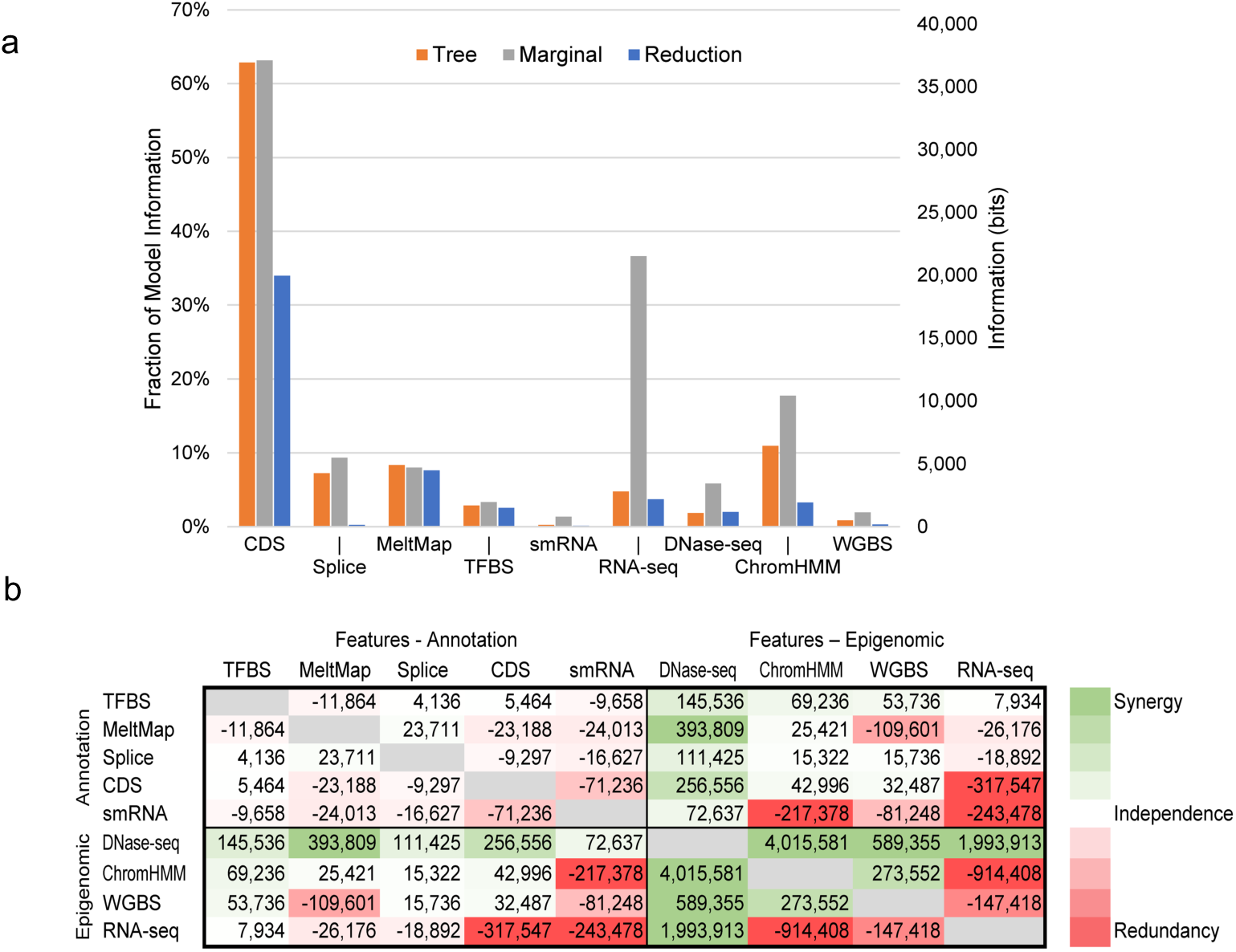
Information and synergy. (a) Information about natural selection is attributed to each individual genomic feature by three methods: (1) summing over the associated decision rules in the tree (tree, orange); (2) measuring the information of the feature in isolation (marginal, gray); and (3) measuring the reduction in total information when that feature is excluded from the complete tree (reduction, blue). These measures are similar when a feature is largely orthogonal to other features (e.g., MeltMap) but different in the presence of strong correlations with other features (e.g., RNA-seq, ChromHMM). (b) Synergy between all pairs of features measured as the excess in information obtained by considering a pair of features together relative to the information obtained by considering the two features separately (Methods). Each cell gives a value in bits. Cells are colored on a spectrum from red (large negative values, indicating redundancy) to green (large positive values, indicating synergy). Note that the matrix is symmetric.

The greedy algorithm used to construct the tree, however, will tend to overestimate the information attributable to features selected early in the process at the cost of features selected later. Therefore, we also considered (1) the *marginal* contribution of each feature in the absence of all others (gray bars in Figure 3a); and (2) the *reduction* in the total information when each feature is removed from the analysis (blue bars). The marginal method attributes 36.6% of all available information to RNA-seq, whereas the reduction method finds that only 3.7% of the available information is specific to RNA-seq. This difference reflects the strong correlation of RNA-seq with CDS annotations. Under the marginal method, the contribution of ChromHMM rises to 17.7% (from 11.0%) and that of DNase-seq to 5.9% (from 1.8%), suggesting substantial correlation of these covariates with one another and/or CDS and RNA-seq. Under the reduction method, the contribution of ChromHMM falls to 3.3%, that of DNase-seq to 2.0%, and that of Splice to 0.3%, with other features being less dramatically affected. Altogether, this analysis shows that the largest share of information about sites that are under selection comes from CDS annotations and RNA-seq data, with ChromHMM coming next and DNase-seq third, but these features are highly correlated with one another. The other annotation types contribute smaller amounts of total information but are less correlated.

### Some genomic features exhibit synergy

The *FitCons2* framework also allows us to ask if there are combinations of features that exhibit *synergy,* in the sense that they yield more total information about natural selection in combination than they do individually. We looked for synergy using a simple pairwise measure defined as the excess in information, in bits, obtained by considering a pair of features together in comparison to the information obtained by considering each feature separately (see Methods). This measure is positive when the combination of two features allows for a better explanation of genome-wide variation as measured by *INSIGHT*, and it is equal to zero when this combination offers no improvement over the individual features (as when the features are nonoverlapping). This measure of synergy can also be negative if two features provide redundant information about how the genome should be partitioned to account for patterns of variation (as when they are strongly correlated along the genome).

We found that most pairs of annotations displayed at most weak synergy (Figure 3b), probably because they tend to identify largely nonoverlapping regions of the genome, and/or to account for few bases overall (as with TFBS, smRNA, and Splice). By contrast, pairs of cell-type-specific epigenomic features often displayed substantial synergy. DNase-seq, in particular, showed synergy with all other epigenomic features, and particularly strong synergy with ChromHMM. In terms of total numbers of bits of synergy, DNase-seq dwarfed all other features. ChromHMM and WGBS also displayed substantial synergy, but at a considerably lower level than DNase-seq. Interestingly, RNA-seq exhibited negative synergy with all epigenomic features except DNase-seq, apparently due to high levels of redundancy.

When pairs of annotations and cell-type-specific epigenomic features were considered, synergy was generally negative or weakly positive, with the exception of DNase-seq, which showed strong positive synergy with all annotations, likely because it signals cell-type-specific activity (see Discussion). In addition, ChromHMM and WGBS each showed weak positive synergy with several annotations, including TFBS and CDS. Altogether, DNase-seq stands out in this analysis as the largest single contributor to synergy, with respect to both annotations and other epigenomic features. ChromHMM and WGBS show some similar trends but to a lesser degree, whereas RNA-seq appears to be the most redundant with other features. These observations have implications for future efforts in data collection and analysis (see Discussion).

### Most entropy derives from mutation and drift

The entropy measured by *FitCons2* reflects a balance of mutation (which acts to increase entropy) with drift and natural selection (which act to reduce entropy)^26-29^. We attempted to separate the contributions of natural selection and the neutral processes of mutation and drift by applying *FitCons2* to a subset of sites assumed to be free from selection for our *INSIGHT* analyses. To allow for heterogeneity across the genome in mutation rates and selection at linked sites, we separately considered such “neutral” sites in each of the 61 clusters identified by *FitCons2* (Methods). By contrasting the entropy per nucleotide site for these neutral sites with the entropy per site for all nucleotides in each cluster, we were able to quantify the reduction in entropy (gain in information) specifically associated with natural selection per cluster.

Across the entire genome, we estimated the neutral entropy per site to be 0.1234 bits, but the actual entropy per site to be 0.1189 bits, indicating a reduction of 0.0045 bits per site from natural selection (Supplementary Table 3). Thus, according to the *INSIGHT* model, natural selection only reduces the entropy in genetic variation that derives from neutral processes by ∼3.6%. However, the relative contributions of neutral processes and natural selection differ considerably by cluster. For example, in cluster 04, which represents splice sites in strongly transcribed coding regions, the neutral entropy per site is estimated at 0.0783 bits, and the observed entropy is 0.0237 bits, a difference of 0.0546 bits. Thus, in this case, most of the available neutral entropy is removed by natural selection. By contrast, in cluster 58, the “NULL” class, the neutral entropy per site is 0.1282 bits and that value is reduced only to 0.1248 bits (a reduction of 2.7%) by natural selection. In general, the reductions in entropy per site due to natural selection are well correlated with estimates of *ρ* (Supplementary Figure 1).

### *FitCons2* scores are informative about the genome-wide distribution of fitness effects, genomic function at individual loci, and relationships among cell types

The average *FitCons2* score per site, across cell types and positions, is 0.082, indicating that an expected ∼8% of nucleotide sites are subject to natural selection, in reasonable agreement with previous measures based on population genetic and phylogenetic data^23,30-33^. These selected sites include an expected 64% of all protein-coding bases (CDS) and 7.6% of all noncoding bases, with more than 90% of sites expected to be under selection falling in noncoding regions. Overall, the highest-scoring positions are in splice sites of annotated genes, followed by protein-coding sequences (CDS), TFBSs, 3’ and 5’ untranslated regions (UTRs), and promoters, with only slight elevations above the background in other annotated elements (Figure 4). These annotation-specific score distributions are often multimodal in a manner that reflects informative combinations of features in the decision tree. For example, the distribution for TFBSs has modes that reflect the partitioning of individual motif positions by information content and the combination with DNase-seq data. Similarly, the UTR and promoter distributions have modes reflecting locally elevated scores, from binding sites, DNase-seq, RNA-seq, WGBS, and related features. Interestingly, the annotation-specific bulk distributions are highly similar across cell types (Supplementary Figure 2), despite considerable differences among the sets of genomic positions they summarize. Thus, while the position-specific *FitCons2* maps are highly cell-type-specific, the differences among cell types are largely explained by cell-type-specific activity.

**Figure 4.**
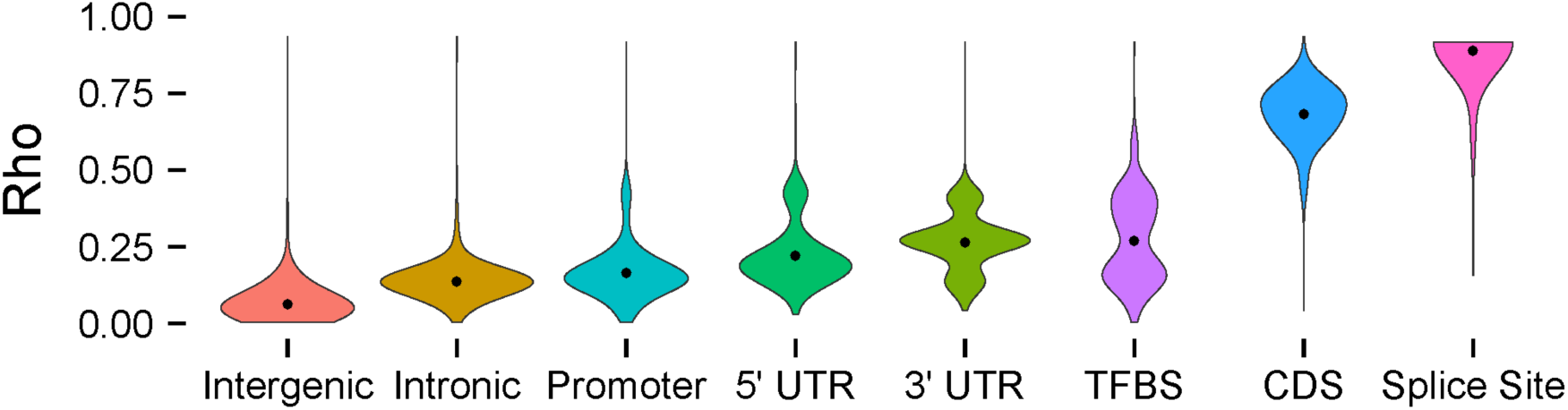
Annotation-specific distributions of *FitCons2* scores. Violin plots showing score distributions for annotated coding regions (CDS), 5’ and 3’ untranslated regions (UTRs), splice sites, transcription factor binding sites (TFBS), core promoters, and remaining intronic and intergenic regions. Scores are for GM12878 cells and reflect regions “active” in that cell type (see Methods). These annotation-specific marginal score distributions are highly similar across cell types, despite differences in regions of the genome they summarize owing to cell-type-specific activity (see Supplementary Figure 2). Splice sites were defined as the two intronic bases immediately adjacent to exon boundaries. Promoters were defined as 1,000 bp upstream of annotated transcription start sites. TFBS annotations based on ENCODE ChIP-seq data were obtained from ref. ^22^.

The *FitCons2* scores across the human genome can be viewed and downloaded via a UCSC Genome Browser track (publicly available at http://compgen.cshl.edu/fitCons2/). This track reveals elevated scores at many enhancers and promoters as well as genes and it often highlights unannotated regulatory elements (Figure 5; see also Supplementary Figures 3 & 4). By zooming into the base level, it is possible to observe high-resolution texture corresponding to features such as individual codon positions, TFBSs, and splice sites (Figure 5a-d). The browser track includes subtracks for the *FitCons2* scores in each of the 115 cell types, which can easily be compared to assess cell-type specificity. In addition, the track includes an “integrated” score that summarizes the scores across all cell types (see Methods) and highlights both cell-type-specific activity and activity shared across cell types (Supplementary Figure 5). This score provides a useful summary when it is not clear to the user what cell type is most relevant in evaluating the functional significance or evolutionary importance of a given site, or when scores are needed for a known cell type that is not among those for which functional genomic data is available.

**Figure 5.**
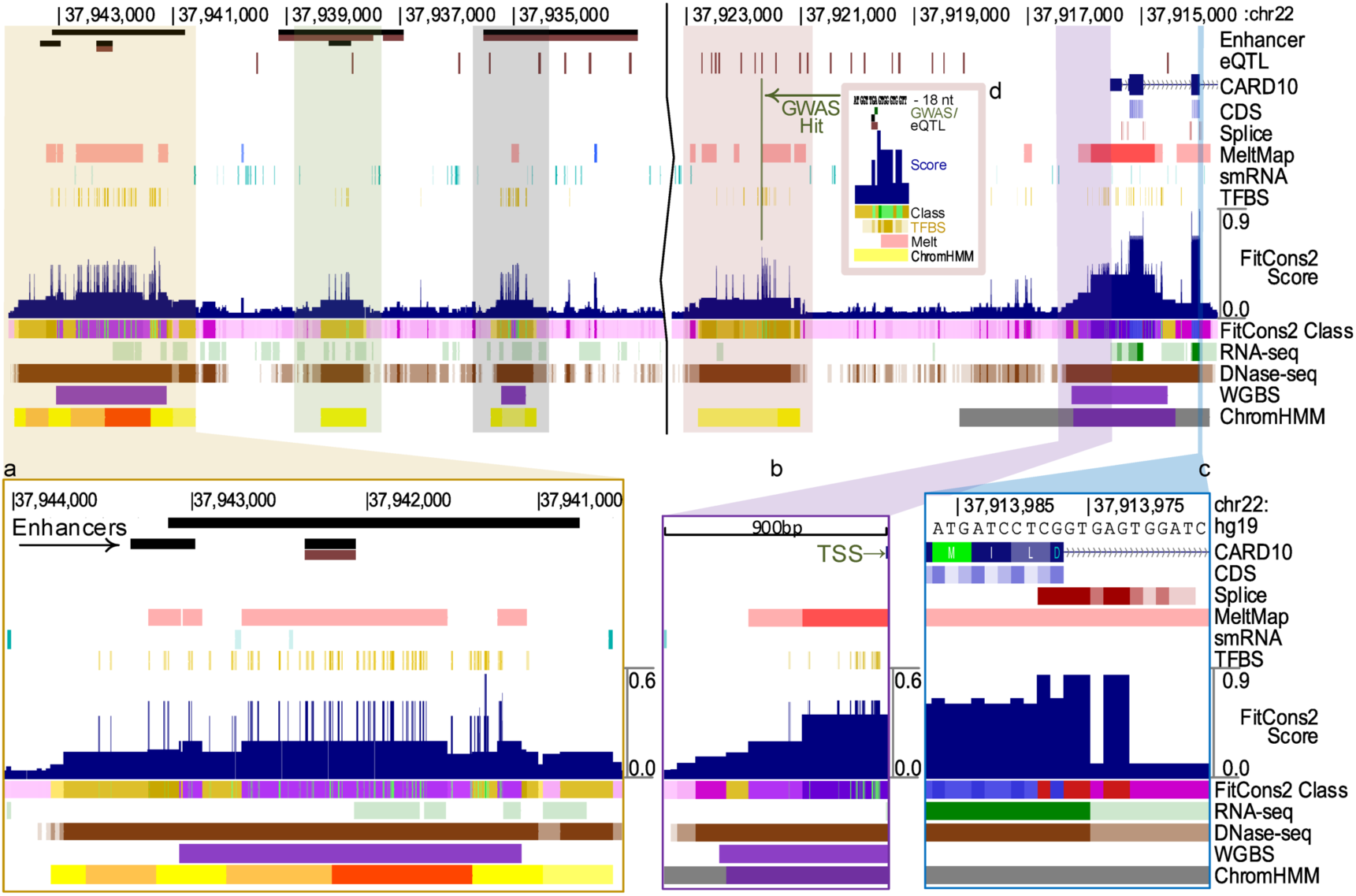
Genome Browser display. UCSC Genome Browser display for a region of chromosome 22 overlapping the 5’ end of the gene encoding caspase recruitment domain-containing protein 10 (CARD10), which participates in apoptosis signaling and activates NF-KB via BCL10. *FitCons2* scores (dark blue, near middle) are shown for the ES-WA7 (embryonic stem cell from blastocyst) cell type. Annotation features are shown above the *FitCons2* scores and cell-type specific epigenomic features are shown at bottom. For reference, predicted enhancers from EnhancerAtlas^51^ (longer bars) and FANTOM5^52,53^ (shorter bars; in both cases, brown indicates enhancers specifically associated with *CARD10*), eQTL from GTEx^54^ (brown indicates specific association with *CARD10*), and the gene annotation from GENCODE^46^ are also shown (top). Insets show zoomed-in displays of (a) an apparent cluster of enhancers showing a high DNase-seq signal, ChromHMM states suggesting regulatory activity (orange: enhancer; red: promoter), and a high concentration of TFBSs; (b) the core promoter and transcription start site showing similar indications of regulatory activity; (c) a 5’ splice site and adjoining CDS and intronic sequences; and (d) GWAS and eQTL hits coinciding with a TFBS. Additional examples are shown in Supplementary Figures 3–5.

While the *FitCons2* scores are primarily designed as an evolutionary measure, we asked whether they are also useful as predictors of genomic function at individual nucleotides. We evaluated the performance of *FitCons2* on two representative prediction tasks: (1) identifying bound TFBSs in a cell-type-specific manner; and (2) distinguishing pathogenic from benign noncoding single-nucleotide variants (SNVs). For comparison, we also considered the original *FitCons*^23^ method (here called *FitCons1*), two well-established measures of phylogenetic conservation (*phyloP*^34^ and *GERP++*^35^), and several other methods that consider both functional and evolutionary genomic information (*CADD*^18^, *LINSIGHT*^24^, *FunSeq2*^20^, and *Eigen*^36^). For the first task, we tested all methods on a high-confidence set of 55,024 predicted binding sites for 12 TFs based on motif matches and supporting cell-type-specific ChIP-seq data from ENCODE^37^, focusing on the H1-hESC and K562 cell-types. On this task, *FitCons2* showed substantially higher sensitivity than all competing methods across score thresholds. For example, at the threshold corresponding to a noncoding coverage equal to the expected fraction of noncoding sites under selection (7.6%; gray vertical line in Figure 6a), the sensitivity for TFBSs of the H1-HESC-specific *FitCons2* scores is 76.8%, compared with 55.2% for *LINSIGHT*, 50.1% for *FunSeq2*, 42.1% for *Eigen*, 39.9% for *FitCons1*, 31.9% for *CADD* and <30% for *GERP*++ and *phyloP*. Results for K562 cells were similar (Supplementary Figure 6). These differences in performance are largely a consequence of *FitCons2*’s unique strategy for making use of information about cell-type-specific “activity” (see Discussion). Nevertheless, the integrated *FitCons2* scores perform roughly as well as the H1-hESC-specific ones, suggesting that our integration scheme is effective in capturing cell-type-specific activity even when cell types differ.

**Figure 6.**
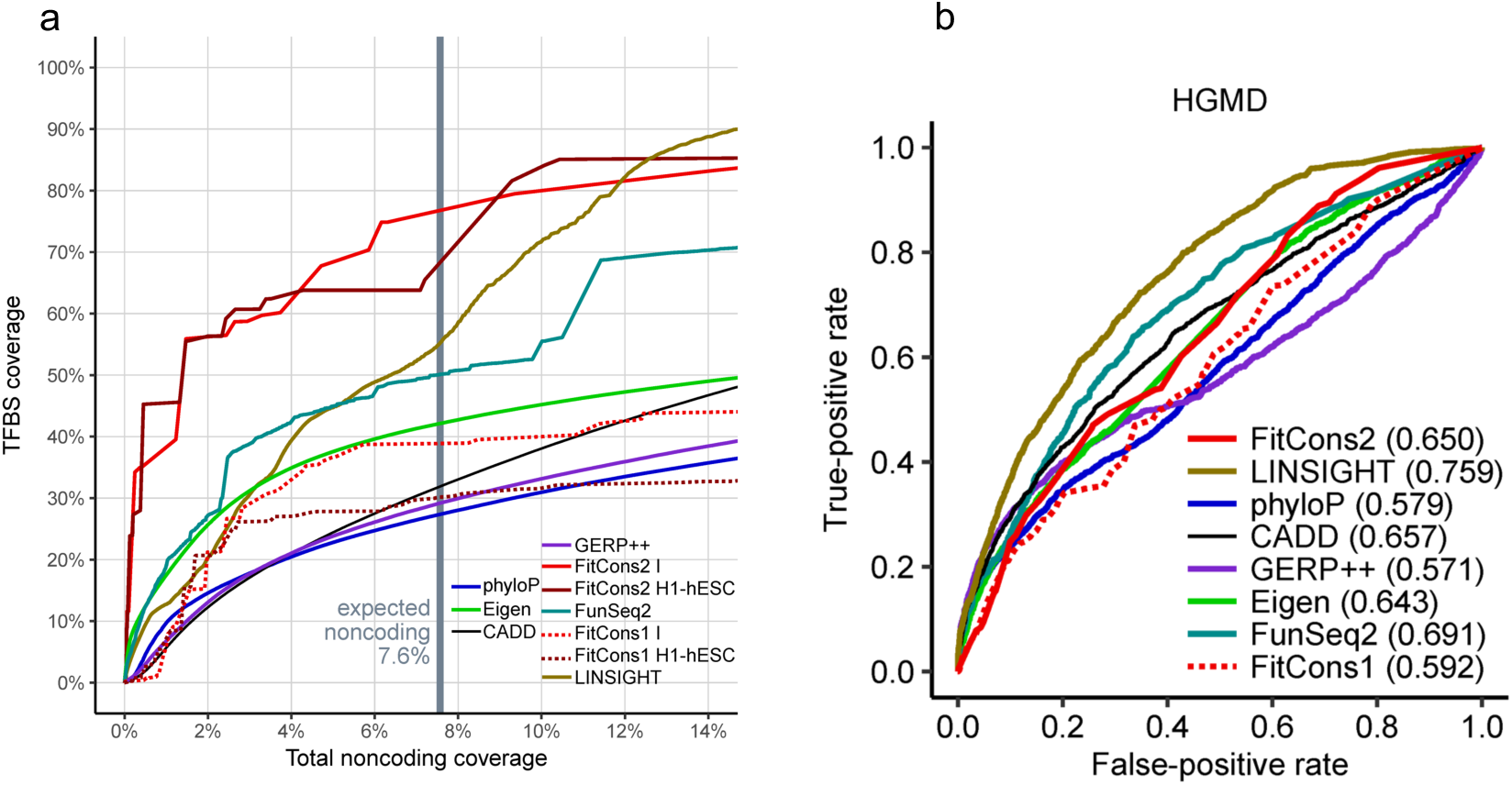
Predictive power for genomic function. (a) Sensitivity of various computational prediction methods (see text) for cell-type-specific transcription factor binding sites (TFBSs). Sensitivity is evaluated using 55,024 motif matches for 12 transcription factors in ChIP-seq peaks for H1-hESC cells^37^ (Methods). Sensitivity is plotted against total coverage outside of annotated coding regions as the prediction threshold for each method is varied. Results for two sets of *FitCons1* and *FitCons2* scores are shown: integrated scores across cell types (I) and cell-type-specific scores for H1-hESC cells. For reference, the vertical gray bar shows the expected fraction of the noncoding genome that is under selection according to *FitCons2* (i.e., the average score in noncoding regions). (b) Receiver operating characteristic (ROC) curves for human disease-associated (pathogenic) single nucleotide variants (SNVs) listed in HGMD. The same computational methods are shown, but in this case only integrated scores are used for *FitCons1* and *FitCons2*. The area-under-the-curve (AUC) statistic is listed after each label in the key. False positives are assessed using likely benign variants matched by distance to the nearest transcription start site (Methods).

For the pathogenic SNVs, we tested the ability of *FitCons2* to distinguish noncoding variants associated with inherited diseases in the Human Gene Mutation Database (HGMD)^38^ from likely benign variants matched by their distance to the nearest transcription start site. In this case, we used our cell-type-integrated scores, because the cell-types of interest vary by disease and in many cases are not known. In this test, *FitCons2* (area under the receiver operating characteristic curve [AUC]=0.650) performs similarly to *CADD* (AUC=0.657) and *Eigen* (AUC=0.643), somewhat better than *FitCons1* (AUC=0.592), *phyloP* (AUC=0.579), and *GERP++* (AUC=0.571), but not as well as *LINSIGHT* (AUC=0.759) or *FunSeq2* (AUC=0.691) (Figure 6b; see also precision-recall curve in Supplementary Figure 7). We also examined variants from the National Center for Biotechnology Information (NCBI) ClinVar database^39^ with similar results, although all methods performed better in this case (and especially the conservation-based methods) owing to an enrichment for splice-site svariants (Supplementary Figure 8). Altogether, we find that *FitCons2* is reasonably competitive with other methods in identifying pathogenic noncoding SNVs, despite its different design goals and its advantages in interpretability and cell-type specificity (see Discussion).

**Figure 7.**
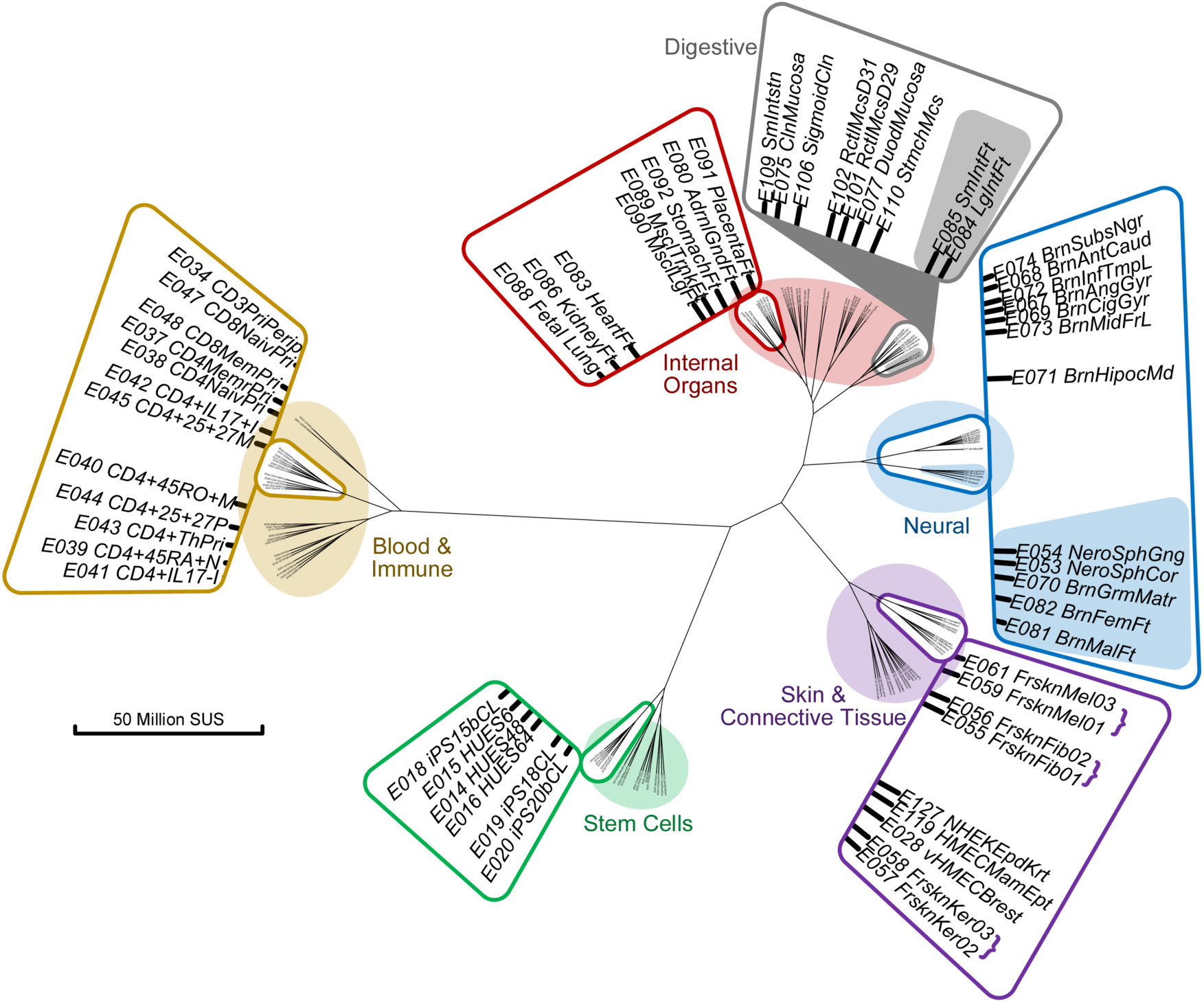
Hierarchical clustering of 115 cell types based on *FitCons2* scores. Dendrogram is derived from a “Manhattan” or *L*1 distance matrix defined such that the distance between every pair of cell types is equal to the sum of the absolute differences of their nucleotide-specific *FitCons2* scores (see Methods). Clustering was done using the default Ward-D2 clustering method in R. Major groups in the dendrogram correspond to cell types associated with (clockwise from top left) blood and the immune system (brown), internal organs (red), the digestive system (gray), neural tissues (blue), skin and connective tissue (purple), and stem cells (green). Insets show examples of closely related cell types from each group. Notice that the digestive cell types are nested within the internal organ-related cell types. Within the neural tissue cluster, separate groups are evident for embryonic and adult brain tissues (blue inset; embryonic cell types highlighted at bottom). Similarly, fetal cell types form subclusters within the internal organ (red inset, entire group) and digestive system (gray inset, gray background) groups. SUS = sites under selection (see text and Methods).

Finally, we asked whether the patterns of cell-type-specific *FitCons2* scores were informative about the relationships among cell types. To address this question, we performed hierarchical clustering of the cell types based on their genome-wide *FitCons2* maps (Figure 7), measuring pairwise distances between cell types in terms of the sum of absolute differences in *FitCons2* scores across all genomic sites. This distance measure is interpretable as the expected total number of nucleotide sites under selection (SUS) based on one cell-type-specific map but not the other (see Methods). This representation naturally groups the Roadmap Epigenomic cell types into ones associated with blood and the immune system, internal organs, the digestive system, neural tissues, skin and connective tissue, and stem cells. Interestingly, induced pluripotent stem cells show similar patterns of genome-wide activity to embryonic stem cells. Brain and neural cells generally cluster together regardless of developmental stage, but within the neural tissue cluster, embryonic brain tissues cluster together and separately from adult brain tissues (see Figure 7 and legend). Similarly, fetal organ and fetal digestive cell-types form sub-clusters within their respective groups.

## DISCUSSION

In this article, we have presented a method for simultaneously clustering genomic sites based on epigenomic features and estimating the probabilities that mutations at those sites will have fitness consequences. Our recursive bi-partitioning algorithm finds clusters of genomic sites that not only share epigenomic features but at which mutations also have similar fitness effects. In statistical terms, our algorithm can be thought of as a model selection procedure that identifies combinations of epigenomic covariates that allow global patterns of genetic variation to be explained by an evolutionary model using the fewest possible free parameters. The algorithm weighs the cost of introducing each new submodel (cluster) against the corresponding benefit in improved model fit. In this way, the procedure resembles stepwise variable selection approaches for regression. Unlike regression methods, however, *FitCons2* is based on a probabilistic evolutionary model and it produces interpretable maximum-likelihood estimates of key evolutionary parameters for each cluster, including, most notably, the probability of fitness consequences, or *FitCons2* score, *ρ*. The interpretability of the *FitCons2* scores—both in terms of the discrete combinations of epigenomic features that define the clusters and the evolutionary meaning of the scores themselves—represents a key advantage in comparison with other available scores for functional relevance or pathogenicity of single nucleotide variants^13-20^. Another major advantage of the scores is that they can be separately computed for each of many cell types to reflect cell-type-specific differences in epigenomic features. We have made available scores for 115 cell types as a UCSC Genome Browser track (http://compgen.cshl.edu/fitCons2/), allowing for convenient visualization, comparison with other genomic annotations, and bulk downloading of data files.

Importantly, this method also allows us to evaluate how informative these features are, both individually and in combination. The individual contributions to information, predictably, are dominated by CDS annotations, with broad, diffuse cell-type-specific epigenomic features such as ChromHMM, RNA-seq, and WGBS, on one hand, and more focused annotations, such as Splice and TFBS, on the other, making smaller but still substantial contributions. The RNA-seq feature stands out as being highly informative by itself but only weakly informative when conditioning on other features, owing to its high degree of redundancy. In our analysis of combinations of features, DNase-seq shows, by far, the most synergy with other features, including both annotations and other cell-type-specific epigenomic features. This synergy appears to result from the ability of DNase-seq to distinguish between “active” and “inactive” elements in a cell-type-specific fashion. For example, the combination of DNase-seq and our cell-type-general annotations of TFBSs provides information about which binding sites are, and are not, occupied in each cell type. When averaging over cell types, this combination further allows for a distinction between TFBSs active in many and those active in few cell types, which are known to experience different levels of constraint^22,24^. This property of DNase-seq suggests that it is a particularly valuable data type to collect in studies in which the budget for functional genomics is limited. Interestingly, while synergy is not directly considered in our recursive bi-partitioning algorithm, the tendency of both DNase-seq and ChromHMM to exhibit synergy with other features appears to be picked up indirectly. For example, at several key nodes (e.g., nodes #2, #3, #11, #12, and #14 in Figure 2), ChromHMM is selected as the most informative feature given previous splits based on CDS, Splice, or MeltMap. DNase-seq has relatively low marginal information so it tends not to get selected early in the recursive algorithm, but it is frequently used near the leaves of the tree (e.g., to distinguish between classes 44 vs. 45 and 28–32 vs. 33–38).

A strength of our method is that it nominally provides cell-type-specific *FitCons2* scores. It is worth emphasizing, however, that the notion of cell-type-specific “fitness” is subtle and potentially misleading. The *FitCons2* scores are ultimately measures of natural selection at the level of whole organisms, based on patterns of genetic variation across a population. The probability that a mutation has fitness consequences at the organism level is not determined separately in each cell type but rather reflects some unknown integral over cell types that depends not only on the importance of the mutation to each cell type but also on the importance of each cell type to the fitness of the organism. Differences across cell types in *FitCons2* scores actually represent differences across cell types in the way sites are grouped by their epigenomic fingerprints, rather than true differences in cell-type-specific fitness consequences. Nevertheless, these cell-type-specific maps are useful in that they effectively capture cell-type-specific activity, allow for sensitive detection of cell-type-specific elements, and reflect the global correlation structure of epigenomic data across cell types. Moreover, our cell-type “integrated” scores attempt appear to provide a useful summary of these scores across all cell types. One way of thinking about these “integrated” scores is that they attempt to approximate the integral across cell types considered by natural selection, making use of our decision tree framework to weight the cell types appropriately (Methods).

The question of how much information is contained in the human genome is not a new one, but that question is typically taken to mean how many bits would be required to encode a single “reference” genome. The answer for the human assembly we have examined here (hg19) is roughly 5.7 billion bits for a simple single-base encoding, or as few as 5.2 billion bits if dependencies between neighboring bases are considered (see Supplementary Text). From an evolutionary perspective, however, this method of measuring information produces a vast overestimate, because most nucleotides in the human genome apparently have no effect on fitness, and in a sense, are not truly “informative” (see ref. ^40^ for a related recent discussion). In addition, human genomes are highly correlated with one another and with the genomes of other primates: given one human genome, or given a chimpanzee genome, another human genome contains much less information that it does alone (roughly 1000th or 100th as much, respectively). Thus, an evolutionary measure of information should consider both fitness effects and population-genetic correlations. With this goal in mind, we use a *population-based* measure of information, and condition on the genome sequences of nonhuman primate outgroups (chimpanzee, orangutan, and rhesus macaque). By making use of a set of putatively neutral sites, we can further decompose the information in a human population into a neutral component (due to a balance between mutation and drift) and component specifically associated with natural selection. Thus, we are able to obtain an approximate measure of the fitness-relevant genetic information in a population of humans, generated and maintained since the human/chimpanzee.

This decomposition reveals that the vast majority of the population-genetic entropy in a collection of human genome sequences, given their primate relatives, can be attributed to a balance between mutation and genetic drift, and that natural selection only slightly diminishes this entropy. This qualitative observation is not surprising, since it is well known that a small minority of nucleotides in the genome are under selection, but it is nevertheless striking that the absolute reduction in entropy, or the information, associated with natural selection is only ∼13 million bits, or ∼1.6 MB—about the size of a typical smartphone snapshot or email attachment. Thus, the fitness-relevant genetic information in a human population, given nonhuman primate genomes, is minimal on the scale of modern digital information, and dramatically smaller than the storage requirements for a single human genome sequence. Our recursive bi-partitioning algorithm can be seen as a way of maximizing the information attributed to selection by making use of heterogeneity along the genome and its epigenomic correlates. However, the genomic features result in an overall information increase of only 58,759 bits (Figure 2), less than half of 1% of the ∼13 million bits associated with selection in the final model. As revealed by the decision tree and our downstream analyses, the information associated with selection is naturally concentrated in coding regions, splice sites, small RNAs, transcription factor binding sites, and other known functional elements, and we find that the *FitCons2* score, *ρ*, is a reasonable surrogate for the degree to which selection contributes to entropy in each of our 61 clusters (Supplementary Figure 1). Nevertheless, it is worth noting that most information still comes from sparsely annotated regions of the genome, because of their enormous size. This apparent information in noncoding regions may be overestimated somewhat in our analysis due to model misspecification, but our estimation method should minimize this effect owing to its reliance on a comparison between neutral and selected regions.

Our approach for measuring evolutionary information bears some resemblance to techniques that make use of an analogy with thermodynamics to quantify an “entropy” associated with natural selection^26-29^. These methods consider the time-dependent distribution of allele frequencies in a population subject to selection and random drift as it approaches equilibrium (stationarity). They define an expectation, called “free fitness” by Iwasa^26^, that is guaranteed to increase to zero as the system approaches its stationary distribution. This quantity is essentially a (negative) relative entropy between the time-dependent and stationary distributions of allele frequencies. Similar to in our analysis, the free fitness can be expressed as a sum of components related to selection, mutation, and drift (see ref. ^29^). However, this framework differs from ours in important respects: for example, the free fitness is an expectation defined with respect to a hypothetical ensemble of independently evolving populations, quantifying it requires knowledge of the population mean fitness, and the emphasis in these papers is on non-equilibrium conditions. Overall, our framework is more heuristic and indirect, but it does allow us to empirically estimate an entropy that results from a balance between mutation and drift by working from a subset of the genome that is putatively free from natural selection, and then to approximate the degree to which selection decreases this entropy by examining the remaining portion of the genome. This reduction of entropy, or information, associated with selection is particularly useful as a relative measure, both for the impact of selection on various genomic regions, and on the relative contributions of selection and neutrality to genomic entropy.

## ACKNOWLEDGMENTS

We thank Ritika Ramani for assistance with browser track development, Noah Dukler for calculating the number of bits required to encode the reference human genome, and other members of the Siepel laboratory for helpful discussions. This research was supported by US National Institutes of Health grants R01-GM102192 and R35-GM127070. The content is solely the responsibility of the authors and does not necessarily represent the official views of the US National Institutes of Health.

## ONLINE METHODS

### Data Sources and Pre-processing

#### Comparative and population genomic data

We measured natural selection using *INSIGHT* and data describing both genetic divergence across primates and polymorphism within human populations. We reused the same data from several previous *INSIGHT*-based analyses^21-24^ (see ref. 22 for complete details). Briefly, these data consist of genome assemblies for chimpanzee (panTro2), orangutan (ponAbe2), and rhesus macaque (rheMac2) aligned to the human reference genome (hg19), together with human polymorphism data extracted from the high-coverage “69 Genomes” data set from Complete Genomics, which was reduced to 54 unrelated samples. Genomic sites were rigorously filtered to eliminate repetitive sequences, recent duplications, CpG sites, and regions not showing conserved synteny across primates. Our analysis considered only the autosomes (chromosomes 1–22) because of substantial differences in mutation rates and distributions of selective effects on the sex chromosomes (X and Y). *INSIGHT* was run using putatively neutral regions identified by starting with all noncoding sites and excluding annotated RNA genes, TFBSs, phastCons-predicted evolutionarily conserved elements, and immediate flanking regions^21,22^. Notably, while much larger population genomic data sets are now available^41-44^, our experiments have shown that the use of even ∼20 times more human individuals makes a negligible difference in estimates of the key parameter *ρ*, owing to the efficiency with which *INSIGHT* pools information across sites in the genome and the property that much of the information about natural selection derives from divergence rather than polymorphism (data not shown). Therefore, we opted to reuse a data set that has already been extensively processed and validated, and whose properties are well known to us.

#### Genomic features

We considered the nine genomic features described in Table 1. For the four epigenomic features, we obtained the imputed RNA-seq, DNase-seq, WGBS, and ChromHMM data sets for each of the 127 cell types (numbered E001–E129, with E060 and E064 omitted) represented in the Roadmap Epigenomic Project data^5^. After initial processing, seven cell types were discarded due to deficiencies in data quality (E001, E003, E017, E027, E098, E104, and E113), and five additional cell types were discarded due to abnormal karyotypes (E114, E115, E117, E118, and E123), which could lead to alignment difficulties and major epigenomic perturbations. For each of the remaining 115 cell types, the “consolidated imputed” RNA-seq and DNase-seq data (representing log RPKM and *p*-values, respectively) were discretized into 4 levels each, using an exhaustive search over possible partition boundaries with an entropy-based objective function (see Supplementary Text for details). The labels from the 25-state version of the Roadmap ChromHMM analysis^5^ were used directly as feature values. The raw WGBS data was partitioned into two classes, corresponding to hypomethylated and non-hypomethylated regions, using the *HMR* program from the *MethPipe* package^45^.

The five annotations were defined as follows for all cell types. The protein-coding gene (CDS) and Splice annotations were derived from the GENCODE V19 database^46^, considering only “KNOWN” “protein_coding” transcripts with a single annotated start and stop codon. Based on CDS annotations, we labeled positions as falling in start codons, codon position 1, codon position 2, codon position 3, and noncoding positions. A position belonging to more than one class across isoforms was assigned to the class under greatest constraint (start > 2 > 1 > 3 > noncoding). For the splice feature, we considered the fifty intronic sites flanking each annotated CDS exon boundary and labeled them, by distance from the exon boundary, as under high, medium, low, or no average constraint, based on pooled data from all splice sites (Supplementary Text). The two positions within CDS immediately adjacent to the exon boundary displayed similar levels of constraint to the “high” intronic class and were included with them. Based on an initial exploratory analysis of potentially relevant genomic features, we also identified predicted DNA melting temperature (MeltMap) as a feature that correlates significantly with selective constraint, although it is likely that this relationship is at least partially explained by the strong correlation of melting temperature with G+C content, which in turn correlates strongly with the presence of functional elements in the genome. In particular, we observed minimal selective pressure at intermediate melting temperatures and elevated selective pressure at more extreme melting temperatures (Supplementary Text). Based on these observations, we discretized the predicted melting temperature into five levels ranging from “very low” to “very high”, with constraint levels such that {very low, very high} > {low, high} > medium.

Because they were available for only a limited collection of cell types, the transcription factor binding site (TFBS) and small RNA-seq (smRNA) features were based on pooled data and treated as annotations. For the TFBSs, we combined 588,958 binding sites from Ensembl Regulatory build V75 (ref. ^47^) with 2,595,018 predicted sites we had previously assembled using ENCODE data^22^. Both sets were derived from ChIP-seq peaks, with bioinformatic post-processing to identify likely TFBSs under the peaks. After merging overlapping predictions, the final set consisted of 1,994,905 TFBSs spanning 23.6Mbp and representing 86 TFs. We partitioned nucleotides into four constraint classes based on the information content of the corresponding position in the position weight matrix for the TF in question (Supplemenary Text). The smRNA data set was based on a combination of the UCSF-4Star composite, the UCSF Brain Germinal Matrix, the UCSC Penis Foreskin Keratinocyte (PFK) composite, and smRNA data from ENCODE for the CD20 and HUVEC cell types. Sites were also partitioned into four levels of constraint based on smRNA data (Supplementary Text).

## Theory and Algorithms

### Recursive bi-partitioning algorithm

The *FitCons2* algorithm begins with the complete set of genomic sites and an associated collection of *D* functional genomic and annotation-based features. Each genomic site is labeled with a particular combination of features, a *D*-dimensional vector known as that site’s functional genomic *fingerprint*. As described above, each of the *D* feature types *i* is discretized into *m*_*i*_ possible values, where *m*_*i*_ ranges between 2 and 25. If these possible values do not have a natural ordering, they are ordered according to their marginal information about natural selection, as measured by the *ρ* parameter from *INSIGHT*. (This ordering by *ρ* is actually performed dynamically at every node in the tree, to allow for changes conditional on previous partitions; see Supplementary Text.) Thus, each nucleotide is assigned one of *k*_*i*_ possible ordered values for each of *D* feature types, *i* ∈ {1, …, *D*}.

The algorithm then considers a family of possible decision rules for splitting the set of genomic sites into two subsets. Each candidate decision rule is based on a single feature type and a threshold. For example, RNA-seq read counts are summarized by four feature values, corresponding to (1) no reads, and (2) low, (3) medium, or (4) high read counts. The algorithm considers partitioning the genome by the decision rules 1*|*234, 12*|*34 and 123*|*4, where *uv | xy* indicates a partitioning between sites labeled *u* or *v* and sites labeled *x* or *y*. Because the feature labels are ordered, the number of possible decision rules for each feature type *i* is always linear in *m*_*i*_. These possible rules must be considered for each of the *D* feature types.

The algorithm selects the decision rule that maximizes the gain in information about natural selection. This choice is made by fitting the *INSIGHT* model separately to the two subsets of genomic sites defined by each candidate decision rule, and deriving a measurement of gain in information from the likelihoods of these models (see below). Choosing partitions that maximize this gain in information has the effect of maximizing the degree to which the resulting two subsets of sites are homogeneous and distinct from one other in terms of their influence from natural selection. Importantly, the gain in information associated with each candidate decision rule is computed as an average over all cell types, that is, by weighting each genomic position by the number of cell types displaying the specified feature value (or range of values) in the *INSIGHT* likelihood function. In this way, the decision tree is fitted to all cell-type-specific data sets simultaneously.

The same procedure is then applied recursively to each of the two subsets of sites, and in turn, to subsets of those subsets, until no subset meets the criterion for further partitioning. Thus, a binary tree is defined with internal nodes representing decision rules and leaves representing particular combinations of decision rules (Figure 1). Furthermore, these leaves define genomic clusters that are maximally homogeneous and distinct in selective pressure. (This greedy algorithm finds a local maximum, but not necessarily a global maximum, according to the objective function used.) Because the algorithm is driven both by the genomic features and the patterns of genetic variation, it tends to find clusters that reflect the natural correlation structure of both the functional genomic and population genomic data.

In practice, we initially had the recursive algorithm terminate when no remaining candidate decision rule provided more than 5 bits of information, which produced a tree with 195 leaves, but then we pruned the tree based on a 50-bit threshold, to obtain a more interpretable 61-leaf tree. This final tree (Figure 1) is unbalanced, with depths ranging from five to twelve. Each step of the recursive algorithm can be viewed as a likelihood ratio test with four degrees of freedom (three free parameters and an addition degree of freedom for the choice of partition), so a 50-bit (69.4-nat) threshold corresponds to a nominal *p*-value of approximately 3 × 10^-14^. Even allowing for the hundreds of tests carried out by the algorithm, this threshold is still conservative. Notice also that, at each step of the algorithm, all internal nodes at a given tree depth can be examined in parallel. Execution of the full algorithm completed in about 57 hours of wall time on a shared computer cluster.

### Algorithm for cell-type integration

To obtain an “integrated” score that summarizes the *FitCons2* scores across cell types, we reused our decision-tree framework to find combinations of cell-type-specific scores that are predictive of overall selective pressure. Our goal was to find a way of summarizing the cell-type specific scores that was not too complex but avoided the pitfalls of a simple average or weighted average — which would tend to be too low in the common case of a site that receives a high score in only one or a few cell types and low scores in many more cell types — or a simple maximum — which would “saturate” quickly and fail to distinguish sites with high scores in many cell types from those with high scores in few cell types.

Briefly, we first summarize the collection of 115 cell-type-specific scores at each position by calculating, at each of 12 different score thresholds between 0 and 1, the number of cell types having scores that exceed that threshold. These numbers of cell types are summarized using five discrete classes per threshold, resulting in a kind of discretized cumulative distribution function at each nucleotide position describing numbers of cell types as a function of score threshold. This calculation is done not with raw counts of cell-types, but using a weighting, based on principal component analysis, that considers the global correlations among cell-type-specific *FitCons2* scores (see Supplementary Text); thus, the CDF can be thought of as describing the number of “effectively independent” cell types exceeding each of the twelve thresholds at each nucleotide site. Our recursive bi-partitioning algorithm is then used to map this 12-dimensional feature vector to a single estimate of ρ per nucleotide site. Specifically, the algorithm clusters nucleotide sites across the genome based on the associated numbers of effectively independent cell types that exceed each designated score threshold, in such a way that the clusters are relatively homogeneous, and relatively distinct from one another, in terms of their influence from natural selection (as measured by *INSIGHT*). Thus, it summarizes the cell-type-specific scores in a manner that reflects the associated global, cell-type-independent patterns of genetic variation. For example, one leaf of the decision tree (one cluster) might represent sites that have at least five (effective) cell types with *ρ* > 0.2; these sites might be assigned an “integrated” estimate of *ρ* = 0.35, based on a direct application of *INSIGHT*. Another leaf of the tree (cluster) might represent sites that have at least ten (effective) cell types with *ρ* > 0.5, and these sites might be assigned an integrated estimate of *ρ* = 0.7. In practice, we use a decision tree with 37 leaves, representing 37 distinct classes.

This approach produces cell-type-integrated scores that summarize the cell-type-specific scores effectively both for sites that are active across many cell types and for sites that are active in a relatively cell-type-specific manner. These integrated scores can not only be used in cases where the desired cell type is not available or not known, but they also provide a good comparison point for non-cell-type-specific scores from other methods. Finally, they provide a useful summary for visualization in the Genome Browser, allowing users to scan the genome for regions of general interest, and then drill down to study the specific patterns of cell-type-specific scores in those regions.

### Measuring Entropy with *INSIGHT*

From an information theoretic perspective, mutation generates entropy in the genome sequences of a population, while genetic drift removes entropy by allowing mutations to become fixed in or lost from the population. Natural selection primarily acts as an additional force for the removal of entropy, by accelerating fixation and loss. (Notably, some forms of selection, such as balancing selection, can act to promote or maintain entropy; we assume such diversifying selection is rare, as it appears to be in humans^48^.) At mutation/selection balance these forces offset one another, and the entropy created by mutation is exactly offset by the removal due to drift and selection^26,29^.

We wish to measure this population genetic entropy from a collection of genome sequences. Moreover, as suggested in the Discussion, we wish to condition on the genome sequences of closely related species and focus on the entropy specific to human populations. Thus, we seek to estimate the entropy of a distribution, *P(X | θ*), where *X* is a collection of human genome sequences and *θ* is a parameter set that governs the distribution (we implicitly condition also on *O*, a collection of closely related nonhuman primate “outgroups”). As shown below, once we can measure this entropy, we can decompose it in various ways, such as by considering contributions from neutral and selective forces, and considering the effects of conditioning on various genomic features.

There is no known algorithm for efficiently characterizing the distribution *P(X | θ*) in a general setting with mutation, recombination, selection, and a complex demographic history, let alone with genomic duplications and rearrangements. Nevertheless, the probabilistic evolutionary model used by *INSIGHT* ^21^ provides an approximate description of *P(X | θ*) that is useful for the purposes of quantifying entropy. Importantly, the *INSIGHT* model is specifically designed to condition on outgroup sequences using a statistical phylogenetic model and to allow for several types of directional selection (including both positive and negative, and both weak and strong, selection). The *INSIGHT* model does not allow for recombination but it does capture aspects of haplotype structure by characterizing a collection of genomes as a series of genomic blocks, each of which has its own mutation and coalescence parameters. In addition, because the *INSIGHT* model describes polymorphisms in terms of allele frequency classes (“low” or “high” derived allele frequencies), it can be thought of as describing the entropy of the entire *population*, rather than of the *sample X* (modulo sampling error in allele frequency estimates).

The *INSIGHT* model is fitted to a collection of genomic sites by maximum likelihood. In the limit of a large number of sites, the maximized log likelihood of the model is closely related to the entropy of the distribution *P(X | θ*), as follows. Conditional on the parameter set, *θ*, and the assumed block structure, *INSIGHT* assumes independence of nucleotide sites, with *P(X | θ*) = *∏*_*i*_ *P(X*_*i*_ *| θ*). Thus, the maximized log likelihood can be written 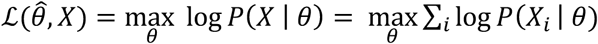. The entropy of 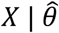, in turn, can be written, 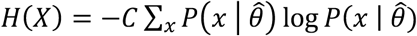, where the sum is over all possible alignment columns *x* and *C* is the number of columns in the actual alignment *X*. Assuming the model fits the data well, in the sense that the distribution of alignment columns under the model is close to the empirical distribution in *X*, then as *C* grows large,

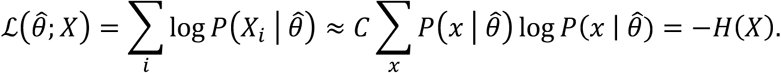

In other words, the negative log likelihood under *INSIGHT* is an estimator for the population genetic entropy. Throughout this article, we assume base-2 logarithms and express entropy in bits.

Note that, in practice, we often compute the log likelihood as an average across cell types, which can be interpreted as the expected complete data log likelihood under a mixture model with a uniform prior. Specifically, for a collection of sites *X*, we assume,

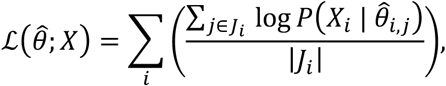

where *j*_*i*_ is the set of cell-types for which data is available at genomic position *i* and 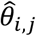 represents the *INSIGHT* model parameters associated with the features in cell-type *j* at position *i*.

### Information associated with features

Suppose a genomic feature *F* allows sites to be partitioned into those having a label (or set of possible labels) *A, X*_*F=A*_ = {*X*_*i*_ *| F(X*_*i*_) = *A*}, and the complement of that set, *X*_*F≠A*_ = {*X*_*i*_ *| F(X*_*i*_) *≠ A*}. A new entropy can be computed based on this partitioning by fitting the *INSIGHT* model separately to *X*_*F=A*_ and *X*_*F≠A*_, with two separate sets of free parameters: 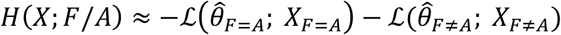. This entropy, *H(X; F/A*), must always be less than or equal to the original entropy, *H(X*) (modulo optimization error). The reason is that the pair of *INSIGHT* models for the two subsets, *X*_*F=A*_ and *X*_*F≠A*_, directly generalizes the single model applied to all sites and must fit the data at least as well, meaning that it will yield a maximized log likelihood at least as large. Thus, 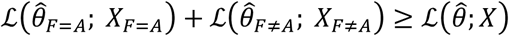, which implies *H(X; F/A*) *≤ H(X*). We can therefore define the nonnegative quantity, 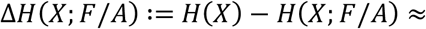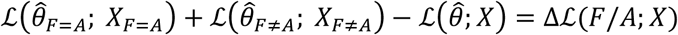, as the “information” associated with feature *F* having label *A*. This is the measure used for the information associated with each decision rule in our recursive bi-partitioning algorithm.

In some cases, it is also useful to have a measure of the overall “marginal” information associated with a feature *F*, considering all of its possible values (e.g., see *Synergy*, below). For this measure, we use:

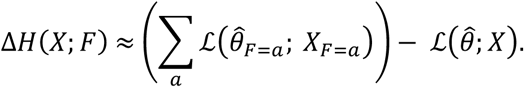

### Partitioning entropy into neutral and selective components

The entropy *H(X*) of a sample of genomic sequences *X* can be described as an additive combination of entropic contributions from mutation, drift, and selection:

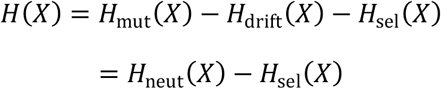

where *H*_neut_(*X*) is the entropy that would exist in the sample if it were evolving free from natural selection, which is itself determined by a balance between the addition of entropy due to mutation, *H*_mut_(*X*), and the subtraction of entropy due to drift, *H*_drift_(*X*); and *H*_sel_(*X*) is the additional entropy removed by (directional) natural selection. In this view, the reduction in entropy represented by *H*_sel_(*X*) can be thought of as the information in *X* that is associated with natural selection. We can similarly write,

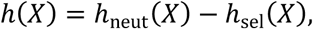

where *h(X*) represents a measure of entropy per nucleotide.

If a subset *X*_*N*_ of alignment columns in *X* can be reasonably assumed to be evolving freely from natural selection, then the neutral entropy per nucleotide, *h*_neut_(*X*), can be estimated using a specialization of the *INSIGHT* model that omits natural selection (i.e., with *ρ* = 0) and describes neutral evolution only:

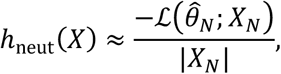

where 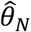 indicates the fitted neutral model and |*X*_*N*_|. is the number of nucleotides in *X*_*N*_. The per-nucleotide reduction in entropy associated with selection can therefore be estimated as,

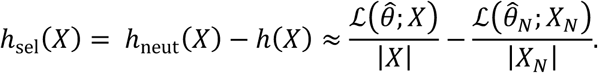

This is a general and interpretable measure of how strongly natural selection has constrained the genetic diversity of the sample. It can be considered a measure of *genetic load* that does not require the estimation of selection coefficients. Notice also that as increasingly rich and precise models of natural selection are considered (as in our recursive bi-partitioning algorithm), *h*_neut_(*X*) will remain constant while *h(X*) decreases, making *h*_sel_(*X*) grow larger and maximizing the information in the sample that can be attributed to natural selection.

In practice, we estimated *h*_sel_(*X*) separately for each of our 61 classes of sites. In order to minimize biases from variation in features such as mutation rate or selection from linked sites, we also estimated *h*_neut_(*X*) separately for each class, using a subset of sites in that class that also fell in our designated putatively neutral sites. In a few cases, this subset was too small to allow the analysis to proceed, so we supplemented the neutral sites in the class with ones located nearby. Our estimates of *h*_sel_(*X*) were generally positive and well correlated with *ρ*, as expected, except for three cases where they turned out to be slightly negative (Supplementary Table 3), apparently because of weak selection and/or sparse data.

## Data Analysis

### Synergy

We define the pairwise synergy between features *F* and *G* as,

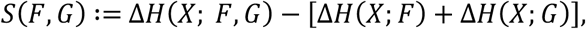

where *ΔH(X; F*) and *ΔH(X; G*) represent the marginal information associated with *F* and *G*, respectively, and *ΔH(X; F, G*) is computed analogously by considering the Cartesian product of feature values for *F* and *G*. *S(F, G*) is positive when there is synergy between *F* and *G*, meaning that more information can be obtained by considering them together than by considering each of them separately; negative when they are “redundant” or highly correlated; and zero when they are, in a sense, orthogonal or “independent”.

In computing *S(F, G*), some special handling is required for sites at which features have a “null” value (meaning a signal that is absent or at background levels). The reason is that there is a kind of trivial “synergy” that derives from sites being removed from the null class of one feature based on a second feature. For example, the null class for feature *F* generally consists of a mixture of true “background” sites and sites that have a positive signal for feature *G*. When *F* and *G* are considered in combination, a component of that mixture is effectively removed from the null class for *F*, making the remaining sites more homogeneous and improving the fit of the model. Thus, a positive “synergy” can be measured even for features with nonoverlapping positive (non-null) values. We therefore exclude all sites at which either *F* or *G* is “null,” which eliminates this trivial contribution to synergy and provides a measure that reflects overlapping positive feature values only.

In most cases, the null feature value is straightforward to define. For example, for epigenomic features such as DNase-seq, RNA-seq, and WGBS, it is the category that represents an absence of any aligned reads, and for annotations such as CDS, TFBS, or smRNA, it generally corresponds to sites at which no annotation is present. In the case of ChromHMM, the null value was taken to be the “quiescent” state, and for MeltMap, it was taken to be the value associated with the weakest selective constraint according to *INSIGHT*.

Notice that *S(F, G)* is similar in spirit to mutual information but conceptually distinct, because it is based on probabilities of a fixed data set *X* conditional on various values of the features *F* and *G*, rather than being based on a probability distribution for *F* and *G*.

### Analysis of predictive power for noncoding genomic function

We examined prediction power for both transcription factor binding sites (TFBSs) and pathogenetic single nucleotide variants (SNVs), limiting our analysis to noncoding regions in both cases. To obtain cell-type specific TFBSs, we obtained the predicted binding sites from ref. ^37^ and intersected them with cell-type-specific ChIP-seq peaks (<500bp in width) from ENCODE. After an examination of five cell-types (GM12878, H1-hESC, He-Las3, HepG2, K562) for which we could identify fairly large numbers of cell-type-specific binding sites for a diverse collection of TFs, we settled on two that were particularly data rich: (1) H1-hESC cells, containing 55,024 TFBSs for 12 TFs (ATF3, CTCF, NANOG, NRF1, NRSF, POU5F1, RFX5, SIX5, SP1, SRF, TCF12, and YY1) spanning 941,645 genomic positions; and (2) K562 cells, containing 66,892 TFBSs for 16 TFs (ATF3, BHLHE40, CEBPB, CTCF, ELF1, NFE2, NRF1, NRSF, RFX5, SIX5, SP1, SRF, TAL1, YY1, ZBTB7A, and ZNF143) spanning 1,080,771 genomic positions. At various score thresholds, we plotted the fraction of nucleotides in TFBS whose *FitCons2* scores exceed the threshold, versus the fraction of all noncoding sites whose *FitCons2* scores exceed the same threshold (Fig 6a and Supplementary Figure 6). This approach produces curves similar to receiver operator characteristic (ROC) curves, but avoids the problem of measuring absolute false positive rates, which is difficult in the presence of incomplete annotations. We plotted analogous curves with scores from *phyloP*^34^, *GERP*++^35^, *CADD*^18^, *LINSIGHT* ^24^, *FunSeq2*^20^, *Eigen*^36^, and *FitCons1*^23^.

For the pathogenic SNVs, we considered noncoding variants associated with inherited diseases in the Human Gene Mutation Database (HGMD)^38^ and the ClinVar database^39^. For each scoring method, we computed false-positive versus true-positive rates for the complete range of score thresholds, displayed the results as ROC and precision-recall curves, and measure prediction power by the area-under-the-curve (AUC or prAUC) statistic. To control for differences in the bulk distributions of scores near and far from genes, we used a matching of HGMD variants with likely benign variants by their distances to the nearest transcription start site that we had previously defined^24^. For *FitCons2*, we based this analysis on the cell-type-integrated scores, because the cell types of interest vary by disease and in many cases are not known.

### Annotation-specific distributions of FitCons2 Scores

The cell-type-specific bulk distributions of scores for various annotation types (Figure 4 and Supplementary Figure 2) were based on regions “active” in each cell type of interest. For annotations associated with protein-coding genes (CDS, splice site, 5’ & 3’ UTR, promoter, and intronic), we defined “active” elements as ones associated with the top third of all annotated genes after ranking them by RPKM based on cell-type-matched RNA-seq data. TFBSs were considered active if they coincided with ChIP-seq peaks in the matched cell type. The notion of cell-type-specific “activity” was not applied to intergenic sites. We took care to exclude any positions that overlapped annotated CDSs from all other categories.

### Hierarchical clustering

The hierarchical clustering of the 115 cell types was based on an *L*_1_ or “Manhattan” distance between each pair of cell types, *C*_1_ and *C*_2_, given by *D(C*_1_, *C*_2_) = *Σi |s*_1,*i*_ - *s*_2,*i*_ *|D*, where *s*_1,*i*_ and *s*_2,*i*_ are the *FitCons2* scores at nucleotide *i* for cell types *C*_1_ and *C*_2_, respectively, and the sum is over all genomic positions. Because *s*_1,*i*_ and *s*_2,*i*_ represent probabilities of natural selection at site *i, D(C*_1_, *C*_2_) can be interpreted as the expected total number of nucleotide sites under selection (SUS) based on one cell-type-specific map but not the other. The 6,555 pairwise distances ranged from 11,433,006 – 61,602,121 SUS. The diagram in Figure 7 was obtained by processing the distance matrix with the default Ward-D2 clustering method in R (V3.3.1).

